# Mu opioid receptor mRNA and protein localization across the rat and mouse habenula

**DOI:** 10.64898/2025.12.16.694766

**Authors:** Aashka K. Popat, Rhiana C. Simon, Beatriz B. Aoyama, Ahana Wokhlu, Aliza T. Ehrlich, Corey C. Harwell, Elyssa B. Margolis

**Author notes:** Functional and Structural Biology Department, State University of Campinas, Campinas, SP, Brazil. Corresponding Author: Elyssa B. Margolis.

## Abstract

The habenula (Hb) has high intensity mu opioid binding and receptor (MOR) expression. It contains medial and lateral subdivisions (MHb and LHb, respectively), yet the details of MOR localization across these regions remains debated. MHb and LHb participate in largely non-overlapping neural circuits, therefore accurately resolving MOR expression across them is critical for understanding how MOR ligands impact behaviors. Here we utilized *in situ* hybridization (ISH) and immunocytochemistry (ICC) to systematically map *Oprm1* mRNA and MOR protein throughout the habenular complex. We studied rat and mouse tissue to evaluate expression across two common research species. We also performed parallel mapping in *Oprm1-^Venus/Venus^* mice. Importantly, we found mRNA and protein in both MHb and LHb in both species. In rat, 39 ± 3% and 21 ± 4% of cells expressed *Oprm1* in MHb and LHb, respectively. These proportions were greater in mouse: 57 ± 1% (MHb) and 32 ± 4% (LHb). *Oprm1* puncta per positive cell were greater in MHb compared to LHb for both rat and mouse (*p* < 0.0001). The highest intensity labeling was localized along the lateral edge of the MHb for all methods. ICC showed MOR localized to fibers and somata in both regions. In LHb, MOR labeling was most dense in intermediate sections along the anterior-posterior (AP) axis. In rats we also observed greater labeling in dorsal LHb at intermediate AP levels and medial LHb more posteriorly. These results indicate that both MHb and LHb can contribute to MOR mediated actions through their respective circuits.

**Key Points:**

- Mu opioid receptor mRNA and protein are expressed in both the medial and lateral habenulae in rat and mouse.
- In the medial habenula, most mu opioid receptor mRNA and protein was detected along its lateral border.
- Across samples, *Oprm1+* cells in the MHb contained more mRNA puncta per cell compared to lateral habenula cells.

## 1 Introduction

The habenula (Hb) is a small, bilateral, evolutionarily conserved structure in the epithalamic region of the diencephalon. It is subdivided into medial and lateral compartments (MHb and LHb, respectively) based on cytoarchitecture and anatomical connectivity (Iwahori et al., 1977; Herkenham and Nauta, 1977; Herkenham and Nauta, 1979). One key feature of the habenular complex is high expression of mu opioid receptors (MORs) (Mansour et al., 1987; Quirion et al., 1983; Recht et al., 1985; Herkenham and Pert, 1980; Gardon et al., 2014). The habenular complex is connected to several forebrain limbic areas and midbrain regions, thereby serving as an important interface for emotional and motivational processing. While both MHb and LHb contribute to encoding aversive signals, they have virtually non-overlapping connectivity and likely have distinct functions (Herkenham and Nauta, 1977; Herkenham and Nauta, 1979). Therefore, an accurate, detailed mapping of the anatomical distributions of MORs across MHb and LHb is critical for developing a framework on which to build an understanding of MOR function in these circuits.

The MHb primarily receives input from the nuclei septofimbrialis and triangularis septum and projects essentially exclusively to the interpeduncular nucleus (IPN) via the fasciculus retroflexus (fr) (Herkenham and Nauta, 1977; Herkenham and Nauta, 1979). The MHb contains both glutamatergic and cholinergic neurons that vary in their patterns of axonal innervation of the IPN (Contestabile et al., 1987; Aizawa, 2012). Activity in this pathway promotes anxiety-like behavior during nicotine withdrawal, and may contribute to processing aversive stimuli and encoding rewarding and negative outcomes (Fowler et al., 2011; Salas et al., 2009; Lee et al., 2019, Handa et al., 2025; Yamaguchi et al., 2013; Sylwestrak et al., 2022; McLaughlin et al., 2017). The MHb–IPN pathway has also been implicated in regulating aversive stress-related behaviors (McLaughlin et al., 2017).

LHb neurons are glutamatergic and receive input from a variety of regions including the ventral pallidum (VP), lateral preoptic area (LPO), entopeduncular nucleus (EPN), lateral hypothalamus (LH), and ventral tegmental area (VTA) (Herkenham and Nauta, 1977; Pasquier et al., 1977; Parent et al., 1981; Skagerberg et al., 1984). Projection targets of LHb include the VTA, substantia nigra pars compacta (SNc), rostromedial tegmental nucleus (RMTg), dorsal and median raphe nuclei (DRN and MRN, respectively), and lateral dorsal tegmentum (LDTg), all via the fr (Herkenham and Nauta, 1979; Cornwall et al., 1990; Christoph et al., 1986; Wang and Aghajanian, 1977; Jhou et al., 2009; Ji and Shepard, 2007; Quina et al., 2015). Increased activity in the LHb produces aversion and depression-like behaviors. For instance, stimulation of glutamatergic inputs to the LHb produces robust aversive behavioral responses, including real-time and conditioned place aversion (Stamatakis and Stuber, 2012; Lammel et al., 2012; Stamatakis et al., 2013; Shabel et al., 2012; Root et al., 2014; Stamatakis et al., 2016; Barker et al., 2017). LHb neurons also increase firing activity in response to rapid aversive stimuli such as foot shock and air puff, and to longer aversive experiences such as immobilization, novel environments, and food deprivation (Benabid and Jeaugey, 1989; Chastrette et al., 1991; Wirtshafter et al., 1994; Timofeeva and Richards, 2001; Lecca et al., 2017). Inactivation via deep brain stimulation and electrolytic lesions of the LHb also reverse depressive-like symptoms (Sartorius and Henn, 2007; Yang et al,. 2008). LHb neurons are excited by negative reward prediction error (RPE) and inhibited by unexpected rewards and positive RPE, thus they encode an inverse RPE, (Matsumoto and Hikosaka, 2007; Matsumoto and Hikosaka, 2009; Hong et al, 2011; Bromberg-Martin, 2011).

As inhibitory GPCRs, MORs that inhibit Hb neurons or their excitatory inputs would be optimally located to dampen neural activity that signals aversive stimuli and behavior states, consistent with MOR agonist induced behavioral effects. Early studies showed that microinjections of morphine into the Hb decrease sensitivity to noxious stimuli and tonic pain (Ma et al., 1992; Cohen and Melzack, 1985). More recently, we showed that microinjecting a MOR agonist into the LHb reverses neuropathic pain-induced nociceptive hypersensitivity and produces affective relief of ongoing pain (Waung et al., 2022). With whole cell electrophysiology, MOR functionality has been detected in both the MHb and LHb. In the MHb, somatodendritic MOR activation decreases spontaneous action potential firing in MOR-expressing MHb neurons; paradoxically, MOR activation on the axon terminals of these neurons in the IPN facilitates glutamatergic transmission (Singhal et al., 2025). MOR activation also somatodendritically inhibits a subset of neurons in the LHb and presynaptically inhibits both glutamate and GABA release onto LHb neurons (Margolis and Fields, 2016; Waung et al., 2022). Together, these studies demonstrate functional MOR localized to both MHb and LHb.

Early anatomical studies used autoradiography with selective MOR ligands to detect binding sites for MOR across the brain; one of the most intensely labeled brain regions in rodents was the Hb (Mansour et al., 1987; Quirion et al., 1983; Recht et al., 1985; Herkenham and Pert, 1980). Subsequent work using specific antisera against the C-terminus of the MOR similarly reported high immunoreactivity in the Hb complex (Arvidsson et al., 1995; Moriwaki et al., 1996). More recent approaches, including an inducible MOR-cre transgenic mouse line and knock-in mouse lines expressing MOR fused to a fluorescent protein, have continued to highlight the density of MOR in the Hb complex (Gardon et al., 2014; Ehrlich et al., 2019; Okunomiya et al., 2020). However, the detailed distribution of MOR across subregions of the Hb remains unclear, and the greatest emphasis has been placed on MOR expression in the MHb in spite of functional evidence for MOR in the LHb. There is also limited information about the expression of *Oprm1*, the gene encoding MORs, in the LHb. Single cell transcriptomic studies of Hb cells suggest that *Oprm1* is more strongly expressed in a subset of MHb neurons compared to other Hb cells (Wallace et al., 2020; Hashikawa et al., 2020), whereas a study looking at the molecular characterization of *Oprm1*-expressing cells using a novel line of *Oprm1*-cre mice reported *Oprm1* cells are found in both MHb and LHb, with a greater number of puncta per cell in the MHb (Mengaziol et al., 2022). To address the lack of consensus and provide greater anatomical detail, here we systematically mapped *Oprm1* mRNA and MOR protein across the rodent Hb with RNAscope *in situ* hybridization (ISH) and immunocytochemistry (ICC), respectively. These approaches enable detection of cells that synthesize MORs and localization of functional MORs. Given the widespread use of transgenic reporter lines to study MOR distribution, we also used *Oprm1*-*^Venus/Venus^* mice (Ehrlich et al., 2019) to characterize MOR-Venus signal across the Hb. This experiment provides information about a widely used tool to compare to our immunocytochemical results, given that each method for localizing MOR protein has strengths and weaknesses. We studied the full AP extent of the Hb, finding mRNA and protein in both MHb and LHb of each species. Resolving these distributions and detailed patterns of MOR expression facilitate improved understanding of possible Hb contributions to opioid mediated behaviors.

## 2 Materials and Methods

### 2.1 Animals

All experiments were performed in accordance with the guidelines of the National Institutes of Health Guide for the Care and Use of Laboratory Animals and approved by the Institutional Animal Care and Use Committees (IACUC) at the University of California San Francisco. Adult male and female C57Bl6/J mice (5-8 weeks old), adult male and female Sprague-Dawley rats (200-300 g), and adult male and female *Oprm1-^Venus/Venus^* mice (8-9 weeks old) (Ehrlich et al., 2019) were used as indicated. All animals were untreated prior to tissue preparation.

### 2.2 Tissue Preparation

Rats were euthanized via intraperitoneal (IP) injection of sodium pentobarbital 200 mg/kg (Euthasol, Virbac) and mice were euthanized with IP 2.5% tribromoethanol in saline (Avertin). All animals were then transcardially perfused with PBS followed by 4% paraformaldehyde, post-fixed in 4% paraformaldehyde at 4°C overnight, and transferred to PBS until sectioning. Brains used for *in situ* hybridization were sectioned within a day of post-fixation to prevent signal degradation; tissue for ICC was sectioned within a week of post-fixation.

Coronal sections (50 μm) containing the Hb were cut using a vibratome (Leica, VT1000S and VT1200S for ICC and RNAscope ISH, respectively). *Oprm1-^Venus/Venus^*tissue was sectioned using a cryostat (Cryostar, NX70). For ISH, sections were collected into a freezing buffer (250 mL total, made with 70 g sucrose, 75 mL ethylene glycol, filled to 250 mL with 0.1 M Na_2_HPO_4_ buffer). Rat brain slices were collected between -2.5 mm and -4.0 mm from bregma (Paxinos and Watson, 2006) and mouse brain slices -1.2 mm to -2.0 mm from bregma (Franklin and Paxinos, 2007). Coordinates referred to in the text were estimated based on atlas landmarks and section thickness.

### 2.3 RNAscope *in situ* hybridization

To measure the expression of *Oprm1* we used the manual RNAscope multiplex platform by ACD Biosystems using the manufacturer’s protocol. We applied DNA probes designed for *Oprm1* (ACD Biosystems #493251 in mice; #410691 in rat) and *Tac2*/*Tac3* (#446391 in mice; #426121 in rats). Images of sections were then captured (40X oil immersion objective) on a confocal microscope (Leica SP8).

### 2.4 RNAscope *in situ* hybridization analysis

Stitched confocal images were organized using Fiji/ImageJ (Schindelin et al., 2012) and imported into QuPath (Bankhead et al., 2017) for analysis. Annotations for MHb and LHb borders were manually drawn using *Tac2*/*3* as a reference for the MHb (Hashikawa et al., 2020). Although there was *Oprm1* signal in the stria medullaris (sm), this region was largely excluded from analysis. Borders between sm and M/LHb differ from ICC tissue in example images due to some slice warping during RNAscope processing. DAPI labeling was used for automated cell detection, and subcellular detection was performed to detect *Oprm1* spots and clusters (Supporting Information Figure 2). We used QuPath’s “estimated spots” output as a proxy for *Oprm1* expression across all cells. Puncta that were counted either overlapped with DAPI signal or the surrounding 5-10 µm around DAPI labeling, indicating that most countied transcripts were likely nuclear. Because we did not observe lateralization in *Oprm1* expression, for fully intact sections where measurements were made on both right and left, the mean of the two sides was taken to produce one value per section. Cells were defined as *Oprm1*+ if they possessed at least 2 spots (puncta). Graphs were produced using custom R scripts and GraphPad Prism. To make statistical comparisons we ran unpaired t-tests on the grand means of *Oprm1+* cell percentage and Welch’s t-test for spot counts because spot variances were different for MHb and LHb (GraphPad Prism 10). For AP position, we ran mixed effects modeling (Type III ANOVA) followed by Tukey’s multiple comparisons testing. Details of statistical tests are included in Supplementary Table 1.

### 2.5 Immunoctyochemistry

Immunocytochemistry was performed on free-floating sections (Aoyama and Popat, 2026; https://dx.doi.org/10.17504/protocols.io.8epv5kejdv1b/v1). Sections were washed with PBS (4x5 min) and then incubated in blocking solution (0.2% bovine serum albumin (BSA), 5% normal goat serum (NGS), and 0.3% Tween20 in S), agitated at RT for 2 hr. Slices were then incubated overnight in rabbit anti-MOR 1:10000 (ImmunoStar Cat# 24216, RRID:AB_572251) (Table 1) with 0.3% Tween20 in PBS, agitated at 4°C. After washes with 0.3% Tween20 in PBS (6x10 min), rat tissue was incubated in 1:500 goat anti-rabbit IgG (H+L) Cy5 (Jackson ImmunoResearch, AB_2338013) (Table 2), and mouse tissue was incubated in 1:500 goat anti-rabbit IgG (H+L) FITC (Jackson ImmunoResearch, AB_2337972) in 0.3% Tween20 in PBS, overnight at 4°C. Tissue was then washed (0.3% Tween20 in PBS, 4x10 min) and incubated (3 hr) at RT in 1:1000 mouse anti-NeuN (Millipore Cat# MAB377, RRID:AB_2298772) (Table 1) in 0.3% Tween20 in PBS, then was washed (0.3% Tween20 in PBS, 4x10 min). Slices were incubated in 1:500 Cy3 Goat anti-mouse (Jackson ImmunoResearch, AB_2338680) (Table 2) for 1 hr agitated at RT. Tissue was washed with 0.3% Tween20 in PBS (4x10 min) then PBS (1x10 min) and mounted using Vectashield mounting medium (Vector, H-1000).

**Table 1:** Primary antibodies used for Immunocytochemistry

| Antibody | Immuogen | Manufacturer, Catalog# | RRID | Host species | concentration |
| --- | --- | --- | --- | --- | --- |
| anti-GFP | Recombinant Full Length Protein corresponding to Aequorea victoria GFP. P42212 | Abcam, ab5450 | AB_304897 | Goat, polyclonal | 1:1000 |
| anti-MOR | Synthetic rat MOR1 (384-398) | Immunostar, 24216 | AB_572251 | Rabbit, polyclonal | 1:10000 |
| anti-NeuN, clone A60 | DNA-binding, neuron-specific protein (NeuN) | EMD Millipore, mab377 | AB_2298772 | Mouse, monoclonal | 1:1000 |

**Table 2:** Secondary antibodies used for Immunocytochemistry

| Conjugate and host | Against | Dilution | Supplier |
| --- | --- | --- | --- |
| Alexa Flour 488 (Donkey) | Goat anti-GFP | 1:2000 | Invitrogen, Cat # A-11055 |
| IgG (H+L) FITC (Goat) | Rabbit anti-MOR | 1:500 | Jackson ImmunoResearch, Cat# 111-095-003 |
| IgG (H+L) Cy5<br>Affinipure (Goat) | Rabbit anti-MOR | 1:500 | Jackson<br>ImmunoResearch,<br>111-175-144 |
| IgG (H+L) Cy3<br>Affinipure (Goat) | Mouse anti-NeuN | 1:500 | Jackson<br>ImmunoResearch,<br>Cat# 111-165-003 |

Venus signal in tissue from *Oprm1-^Venus/Venus^* mice (Ehrlich et al., 2019) was amplified with immunocytochemistry using an anti-GFP antibody (Table 1). Tissue was permeabilized with 0.2% Triton-X-100 in PBS (1x10 min) and then incubated in blocking solution for 30 mins (3% BSA, 0.3% Triton-X100 in PBS), agitated at RT. Tissue was then incubated at 4°C overnight in goat anti-GFP 1:1000 (Abcam Cat# ab5450, RRID:AB_304897) diluted in blocking solution.

After washes in 0.2% Triton-X-100 in PBS (2x10 min), sections were incubated in 1:2000 AF488 donkey anti-goat (Invitrogen, Cat # A-11055, RRID: AB_2534102) (Table 2) diluted in blocking solution for 1hr at RT. All tissue was washed with 1:10000 Hoechst-33342 in PBS (Thermo Scientific, cat# H1399) as a counterstain, washed in PBS (2x10 min), and mounted with mounting medium (HelloBio, cat# HB9854). This tissue was also stained with a mouse anti-NeuN monoclonal antibody (EnCor Biotechnology Cat# MCA-1B7, RRID:AB_2572267) and an AF594 donkey anti-mouse secondary (Thermo Fisher Scientific Cat# A-21203, RRID:AB_2535789), but Hoechst-33342 showed a more reliable signal for identifying Hb regions in this tissue.

Immunolabeled sections were imaged using a confocal LSM (Olympus FV4000, Fluoview). Stitched z-stacks of the bilateral habenula were captured with a 20x objective for mouse tissue and 40x objective for rat tissue. Example micrographs for each were captured with a 40x objective in z-stacks.

### 2.6 Immunocytochemistry analysis

To register images for averaging signal intensity across animals, the border between MHb and LHb was identified independent of MOR labeling. Images were aligned across slices using Big Warp on maximal projections of z-stack tile scans (Bogovic et al., 2016, Pietzsch et al., 2015) in Fiji (Schindelin et al., 2012). Density of NeuN labeling was used to identify the M-L border except in *Oprm1-^Venus/Venus^*tissue where Hoechst-3342 was used (Iwahori, 1977). Landmarks used for aligning samples were agreed upon by two researchers to reduce bias.

The MOR channel data from registered images were exported as pixel intensity matrices that were averaged and normalized across animals to create heatmaps of MOR expression at each AP coordinate. Since we did not find any lateralization in MOR expression, values for left and right hemispheres were averaged together. Slices were excluded from registration and analysis if there were ruptures in the tissue that prevented accurate registration or if the section was not within 100 μm of the target coordinate.

### 2.7 Antibody Characterization

The rabbit anti-MOR antibody (ImmunoStar Cat# 24216, RRID:AB_572251) targets the C-terminus of MOR (AA 384-398). The manufacturer reports that preadsorption with MOR peptide (384–398) at 10 µg/ml completely eliminates labeling. The specificity of the antiserum has also been established in studies using immunolabeling of transfected cells and immunoisolation (Arvidsson et al., 1995). We observed that immunocytochemistry performed with this antibody in the dorsal striatum of a MOR-KO mouse (*Oprm1^tm1Kff^*; RRID:IMSR_JAX:007559; Matthes et al., 1996) produced no signal (Supporting Information Figure 1).

**Figure 1.**
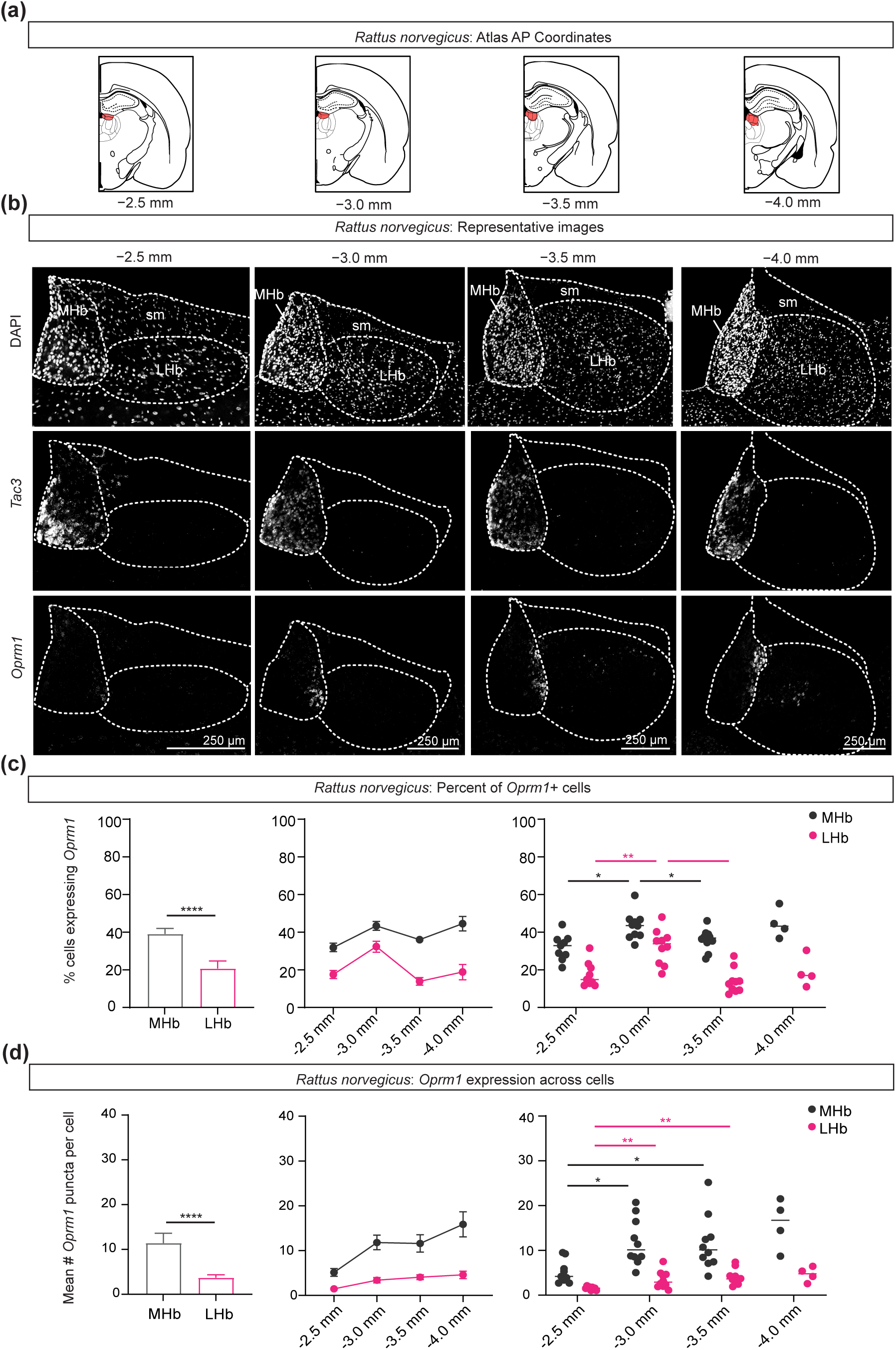
*In situ* hybridization for *Oprm1* in *Rattus norvegicus* (n = 5 male, 5 female Sprague-Dawley rats). **(a)** Depictions of coronal sections at four AP levels chosen for processing (∼-2.5 mm, -3.0 mm, -3.5 mm, -4.0 mm from bregma). The habenular complex is highlighted in red. Coordinates and drawings are based on the rat brain atlas (Paxinos and Watson, 2006). **(b)** Representative images for DAPI (top), *Tac3* RNAscope (middle), and *Oprm1* RNAscope (bottom) across the habenula for one rat. Scale bar = 250 μm. **(c)** Percent of cells expressing *Oprm1* in the MHb and LHb across the AP axis (N = 10). Grand mean across all sections and rats (left; unpaired t-test), average percent for each section (middle), and values by individual rats (right; Type III ANOVA for habenular subregion x AP position, Tukey’s multiple comparisons test). Sections within ∼250 μm were binned together. A cell was defined as *Oprm1*+ if it possessed at least 2 puncta. **(d)** Mean number of *Oprm1* puncta per cell across all *Oprm1*+ cells in the MHb versus the LHb. Grand mean across all sections and rats (left; Welch’s t-test), average number of puncta/cell for each section (middle), and values separated by individual rats (right; Type III ANOVA for habenular subregion x AP position; Tukey’s multiple comparisons test). Lhb, lateral habenula; MHb, medial habenula; sm, stria medullaris. * *p* ≤ 0.05, ** *p* ≤ 0.01, **** *p* ≤ 0.0001.

The mouse anti-NeuN antibody (Millipore Cat# MAB377, RRID:AB_2298772) targets the DNA-binding, neuron-specific protein (NeuN/RBFOX3). As specified by the manufacturer, this antibody recognizes 2-3 bands in the 46-48 kDa range and possibly another band at 66 kDa. Signal is detected in most CNS and PNS neuronal cell types of all vertebrates tested, and signal was not present in non-neuronal tissue such as fibroblasts (Mullen et al., 1992).

The goat anti-GFP antibody (Abcam Cat# ab5450, RRID:AB_304897) recognizes the recombinant full length protein corresponding to Aequorea GFP.P42212. This antibody has been verified in Western Blot samples labeling a band at 27 kDa (manufacturer specified).

For secondary antibodies, we observed some non-specific goat anti-rabbit IgG (H+L) Cy5 (Jackson ImmunoResearch, AB_2338013) binding in mice and therefore used goat anti-rabbit IgG (H+L) FITC (Jackson ImmunoResearch, AB_2337972) in mouse tissue, which greatly reduced background fluorescence. We used the hippocampal dentate gyrus layer just dorsal to the Hb as a control region to evaluate non-specific binding.

## 3 Results

We mapped *Oprm1* mRNA and MOR protein across the AP extent of the Hb in adult male and female SD rat and C57Bl6/J mouse. ISH (RNAscope) and ICC were performed in tissue from separate animals. We also compared these distributions to that in *Oprm1-^Venus/Venus^*mice.

### 3.1 *Oprm1* distribution in rat Hb

We used RNAscope to label *Oprm1* mRNA of Sprague-Dawley rats (Figure 1; N = 5 female, 5 male). In order to identify the border between MHb and LHb accurately, we used both DAPI labeling and RNAscope against *Tac3* mRNA, an effective marker for MHb (Hashikawa et al., 2020). In this approach, each puncta represents a single mRNA molecule, and *Oprm1*+ cells were defined as those possessing at least two puncta. One-punctate cells were excluded due to the possibility of noise, but given the possibility of cells not requiring many *Oprm1* transcripts in order to produce adequate amounts of protein, we counted low *Oprm1*-expressing cells as positive (2 or more puncta). *Oprm1* signal from the sm and fr were excluded from quantification.

In the MHb, cells with a high-intensity *Oprm1* signal were predominantly localized to its most lateral aspect (Figure 1b). These *Oprm1*+ cells were more apparent in intermediate AP slices and signal intensity appeared weakest in anterior MHb, at -2.5 mm posterior to bregma (Figure 1b). *Oprm1*+ cells that had fewer puncta were present more medially, in the dorsal MHb (Figure 1b). In the LHb, we observed puncta scattered throughout all sections, with overall fewer puncta per cell compared to MHb (Figure 1b,d). We noted that there were cells in posterior slices that expressed more intense *Oprm1* signal. For instance, at -3.5 mm posterior to bregma, distinct mRNA signal was present in the dorsolateral LHb (Figure 1b). At -4.0 mm posterior to bregma, a few *Oprm1*+ cells with higher mRNA expression were located in the medial LHb (Figure 1b).

*Oprm1* signal in these representative images reflected quantitative patterns of expression. On average, the percentage of *Oprm1+* cells in the MHb was greater than that in the LHb (*p* < 0.0001): 39 ± 3% of MHb and 21 ± 4% of LHb cells were *Oprm1*+ (Figure 1c). These percentages also varied across AP position (*p* = 0.002; Figure 1c). In both the MHb and the LHb percentages of *Oprm1+* cells were significantly greater -3.0 mm posterior to bregma compared to -2.5 mm (*p* = 0.03 for MHb; *p* = 0.009 for LHb) and -3.5 mm (*p* = 0.03 for MHb, *p* = 0.0004 for LHb) posterior to bregma (Figure 1c). *Oprm1+* cells possessed more puncta per cell in the MHb compared to the LHb, averaging 11 ± 2 puncta/cell in the MHb and 4 ± 1 puncta/cell in the LHb (*p* < 0.0001; Figure 1d). Average mRNA expression also varied by AP position (*p* = 0.001); *Oprm1+* cells at -2.5 mm from bregma possessed fewer puncta/cell than at -3.0 mm (*p* = 0.02 MHb; *p* = 0.03 LHb) and at -3.5 mm (*p* = 0.04 MHb; *p* = 0.003 LHb) from bregma (Figure 1d).

### 3.2 *Oprm1* distribution in mouse Hb

We labeled *Oprm1* and *Tac2* mRNA across the Hb (Figure 2) in C57Bl6/J mice (N = 6 female, 6 male). As above, borders were defined using a DAPI counterstain density and *Tac2* as a marker for MHb (Hashikawa et al., 2020). Signals from the sm and fr were excluded from quantification.

**Figure 2.**
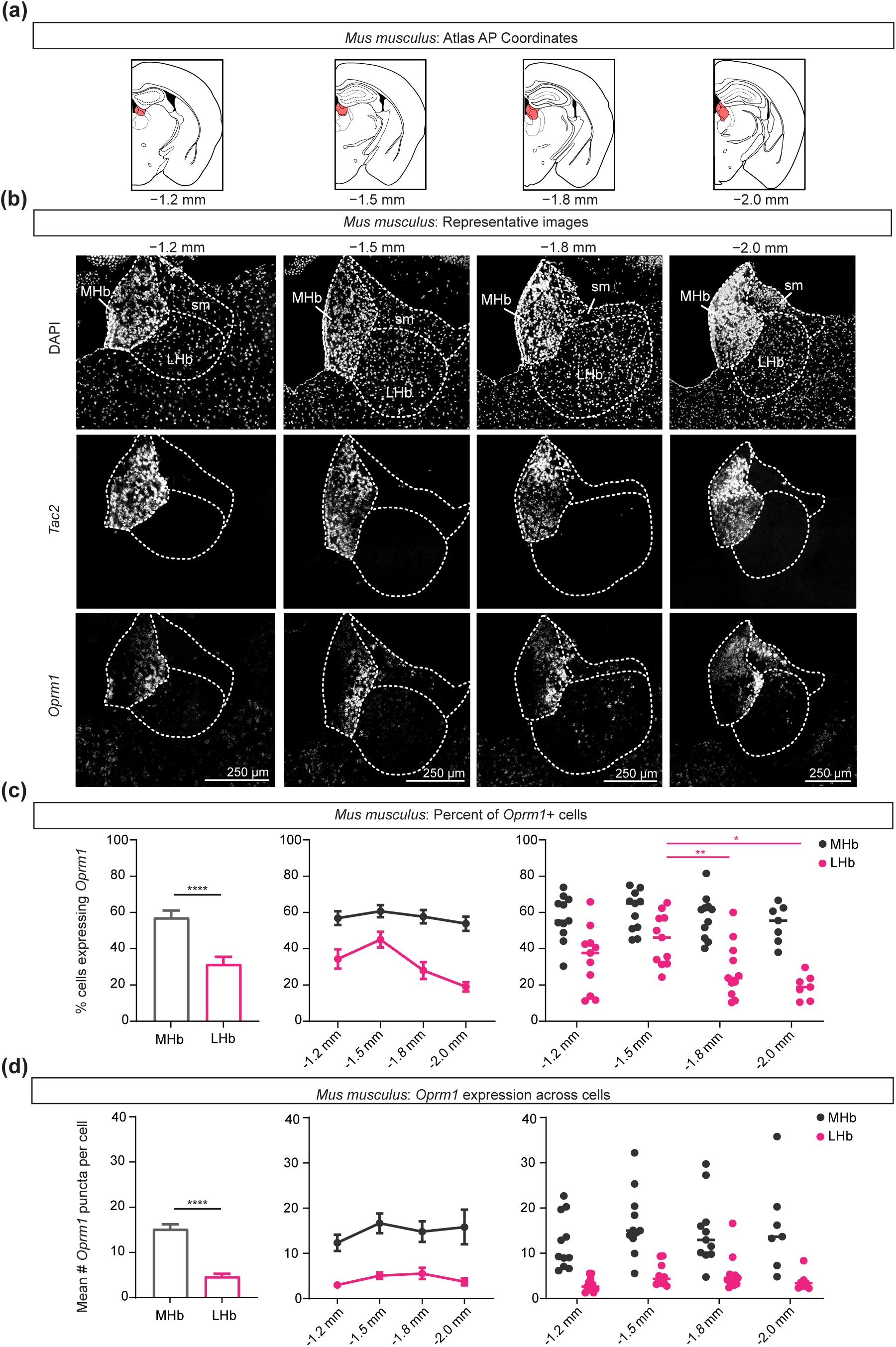
*In situ* hybridization for *Oprm1* in *Mus musculus* (n = 6 male, 6 female C57Bl6/J mice). **(a)** Depictions of coronal sections at the four AP levels chosen for processing (∼-1.2 mm, -1.5 mm, -1.8 mm, - 2.0 mm from bregma). The habenular complex is highlighted in red. Coordinates and drawings are based on the mouse brain atlas (Franklin and Paxinos, 2007). **(b)** Representative images for DAPI (top), *Tac2* RNAscope (middle), and *Oprm1* RNAscope (bottom) across the habenula for one mouse. Scale bar = 250 μm. **(c)** Percent of cells expressing *Oprm1* in the MHb and the LHb across the AP axis. Grand mean across all sections and mice (left) (unpaired t-test), average percent for each section (middle), and values separated by individual mice (right; Type III ANOVA for habenular subregion x AP position, Tukey’s multiple comparisons test). Sections within ∼100 μm were binned together. A cell was defined as *Oprm1*+ if it possessed at least 2 puncta. **(d)** Mean number of *Oprm1* puncta per cell across all *Oprm1*+ cells in the MHb and LHb. Grand mean across all sections and mice (left), average number of puncta/cell for each section (middle; Welch’s t-test), and values separated by individual mice (right). Lhb, lateral habenula; MHb, medial habenula; sm, stria medullaris. * p ≤ 0.05, ** p ≤ 0.01, **** p ≤ 0.0001.

We observed high intensity *Oprm1* labeling in cells along the lateral edge of the MHb (Figure 2b). This high intensity signal was present at all AP levels including the most anterior section, unlike rat, and occupied a wider ML extent of MHb (Figure 1b,2b). There were also puncta present throughout the LHb, with moderate intensity signal distributed differently along the AP axis (Figure 2b). In more detail, at -1.5 mm posterior to bregma there was bright *Oprm1* labeling in the medial LHb, close to the MHb border, and in the dorsal LHb (Figure 2b). At -1.8 mm posterior to bregma, there was clear signal from the lateral half of the LHb (Figure 2b). In the most posterior LHb (-2.0 mm from bregma), there were *Oprm1*+ cells with higher numbers of puncta in the dorsal and medial LHb, similar to rat (Figure 1b, 2b).

As in rat, in mouse the average percentage of cells labeled for *Oprm1* was higher in the MHb (57 ± 1%) compared to LHb (4 ± 1%; *p* < 0.0001; Figure 2c). Average percentages were not statistically different across AP position in the MHb, while they were greater in the intermediate LHb at -1.5 mm (*p* = 0.005) and -1.8 mm (*p* = 0.01) compared to posterior LHb (-2.0 mm from bregma; Figure 2c). On average, *Oprm1+* cells in the MHb possessed more puncta/cell than in the LHb: we detected 15 ± 1 puncta/cell in MHb and 5 ± 1 puncta/cell in LHb (*p* < 0.0001; Figure 2d). These values did not differ across AP positions in the MHb or LHb (Figure 2d).

### 3.3 MOR distribution in the rat habenula

We performed ICC to localize MOR protein in the Hb of Sprague-Dawley rats (N = 3 female, 3 male). Tissue was counterstained with NeuN, and neuronal density was used to identify the border between M- and LHb (Iwahori, 1977). NeuN distributions were comparable to DAPI (Figure 1b), with the exception of less NeuN labeling in the sm compared to DAPI labeling, consistent with the expectation of sparser neuronal cell bodies in this fiber track.

MOR signal was evident throughout the AP extent of the MHb (Figure 3). At the most anterior level, there was sparse labeling along the ventral part of the medial edge of the MHb-LHb border (Figure 3b,c_i_). This signal was greater at intermediate AP levels and shifted dorsally in more posterior slices (Figure 3b,c_i_). Labeling of MOR protein in this region of the MHb appeared localized to both somata and fibers (Figure 3c_i_).

**Figure 3.**
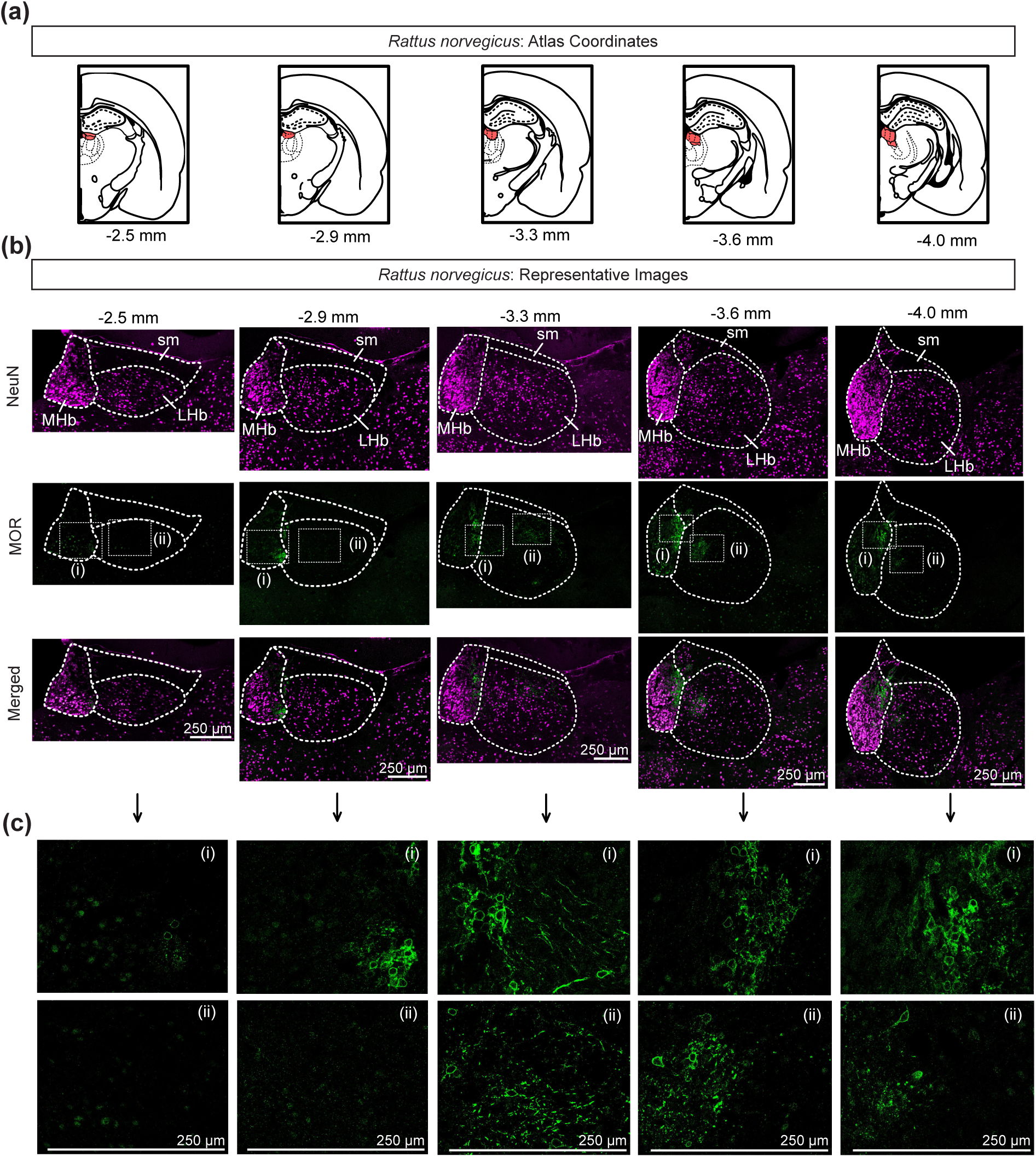
Immunocytochemistry for MOR protein in *Rattus norvegicus* (n = 3 male, 3 female Sprague-Dawley rats). **(a)** Depictions of coronal sections at the five AP levels chosen for immunocytochemical processing (∼-2.5 mm, -2.9 mm, -3.3 mm, -3.6 mm, -4.0 mm from bregma). The habenular complex is highlighted in red. Coordinates and drawings are based on the rat brain atlas (Paxinos and Watson, 2006). **(b)** Representative images of NeuN ICC (magenta, top row), MOR ICC (green, middle row), and merged channels (bottom row) were chosen from one rat for each AP coordinate. **(c)** Representative images of MOR ICC in MHb (i) and LHb (ii) corresponding to the boxed areas in **(b)**. Images in **(b)** and **(c)** depict one focal plane of their respective z-stacks. Scale bars = 250 μm. LHb, lateral habenula; MHb, medial habenula; sm, stria medullaris.

MOR expression was sparse in the anterior LHb, with higher intensity signal in intermediate and posterior sections (Figure 3b, c_ii_). For instance, there was strong immunoreactivity at -3.3 mm posterior to bregma across the LHb in both the medial and dorsolateral LHb (Figure 3b). In the medial LHb, MOR protein seemed localized primarily to fibers (Figure 3b,c_ii_). In the dorsolateral quadrant of the LHb, we observed MOR-labeled fibers and cell bodies (Figure 3c_ii_). In posterior sections (-3.6 mm, -4.0 mm from bregma), MOR labeling in the LHb was brightest medially, including more outlined cell bodies, compared to medial LHb MOR labeling -3.3 mm from bregma (Figure 3b,c_ii_).

These patterns of MHb and LHb labeling were consistent across rat samples. When aligned and averaged across animals, there was a clear intensity in a thin strip of MOR labeling in the lateral MHb at the border between the two regions (Figure 5a). This density expanded dorsally in more posterior sections (Figure 5a). In the LHb, MOR was consistently detected at more moderate levels in the medial LHb and throughout the dorsal half of the lateral LHb. At -3.6 mm posterior to bregma, there was notable labeling in the medial LHb (Figure 5a). A more confined, medial area was consistently labeled at -4.0 mm posterior to bregma (Figure 5a).

### 3.4 MOR distribution in mouse Hb

We mapped MOR protein with ICC in the Hb of C57Bl6/J mice (N = 3 female, 2 male). The topography of NeuN signal in the Hb looked similar between rat and mouse tissue throughout the AP axis (Figure 3b, 4b).

There was dense MOR labeling along the lateral edge of MHb throughout the AP extent of the Hb (Figure 4b). Higher magnification shows most of this labeling outlining somata (Figure 4c_i_). In contrast to rat where signal in this area shifted across AP position, in mouse it spanned the dorsoventral extent of the MHb along the entire AP axis (Figure 3b,4b,5).

**Figure 4.**
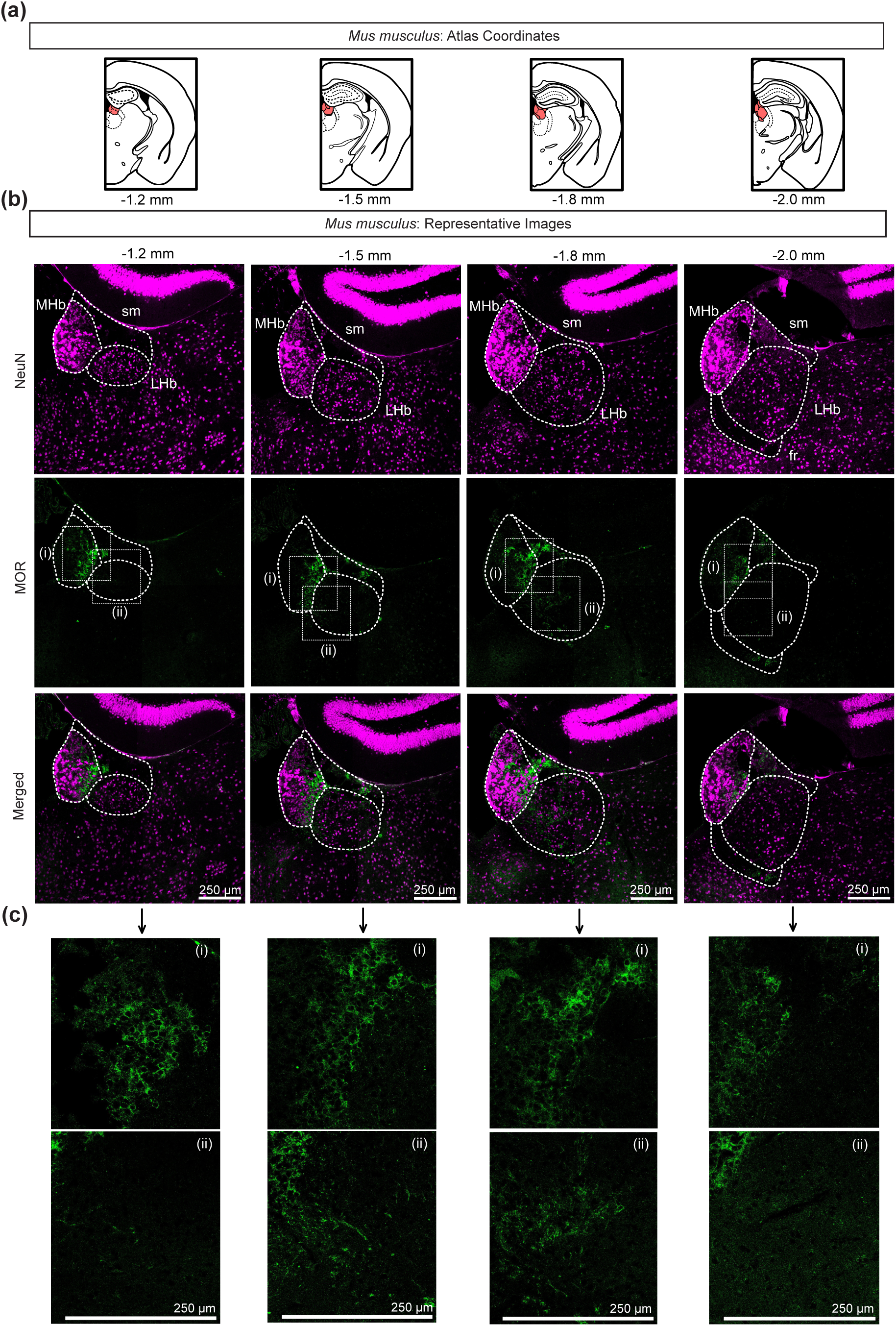
Immunocytochemistry for MOR protein in *Mus musculus* (n = 2 male, 3 female C57Bl6/J mice). **(a)** Depictions of coronal sections at the four AP levels chosen for immunocytochemical processing (∼-1.2 mm, -1.5 mm, -1.8 mm, -2.0 mm from bregma). The habenular complex is highlighted in red. Coordinates and drawings are based on the mouse brain atlas (Franklin and Paxinos, 2007). **(b)** Representative images of NeuN ICC (magenta, top row), MOR ICC (green, middle row), and merged signal (bottom row) were chosen from one mouse for each AP coordinate. **(c)** Enlarged images of MHb (i) and LHb (ii) in **(b)**. Images in **(b)** and **(c)** depict one focal plane of their respective z-stacks, and (i) and (ii) depict different focal depths. Scale bars = 250 μm. LHb, lateral habenula; MHb, medial habenula; sm, stria medullaris; fr, fasciculus retroflexus.

In the LHb, in anterior sections (-1.2 mm from bregma) we mostly detected labeling in fibers, particularly in the ventromedial LHb (Figure 4c_ii_). This signal was also present at -1.5 mm posterior to bregma, with the addition of a few interspersed outlined cell bodies (Figure 4c_ii_). There was a small cluster of fibers and outlined cell bodies in the laterodorsal LHb at -1.5 mm from bregma, close to but distinct from labeling in the sm (Figure 4b). At -1.8 mm, we observed the most extensive MOR signal, primarily in the medial half of the LHb (Figure 4b,c_ii_). MOR localization in this area consisted of several fibers and some outlined somata towards the lateral extent of the medial LHb (Figure 4c_ii_). MOR signal in the LHb was sparse at the most posterior level of the LHb (Figure 4b,c,5).

Across mice we observed a layer of intense MOR labeling in the MHb along most of the border with LHb (Figure 5b). Heatmaps of MOR distributions averaged across animals also show clear labeling in the LHb in intermediate sections (-1.5 mm, -1.8 mm posterior to bregma), with somewhat higher intensity in the medial LHb (Figure 5b). There was also distinct MOR labeling in the sm in the mouse tissue that was not clearly evident in rat (Figure 5).

**Figure 5.**
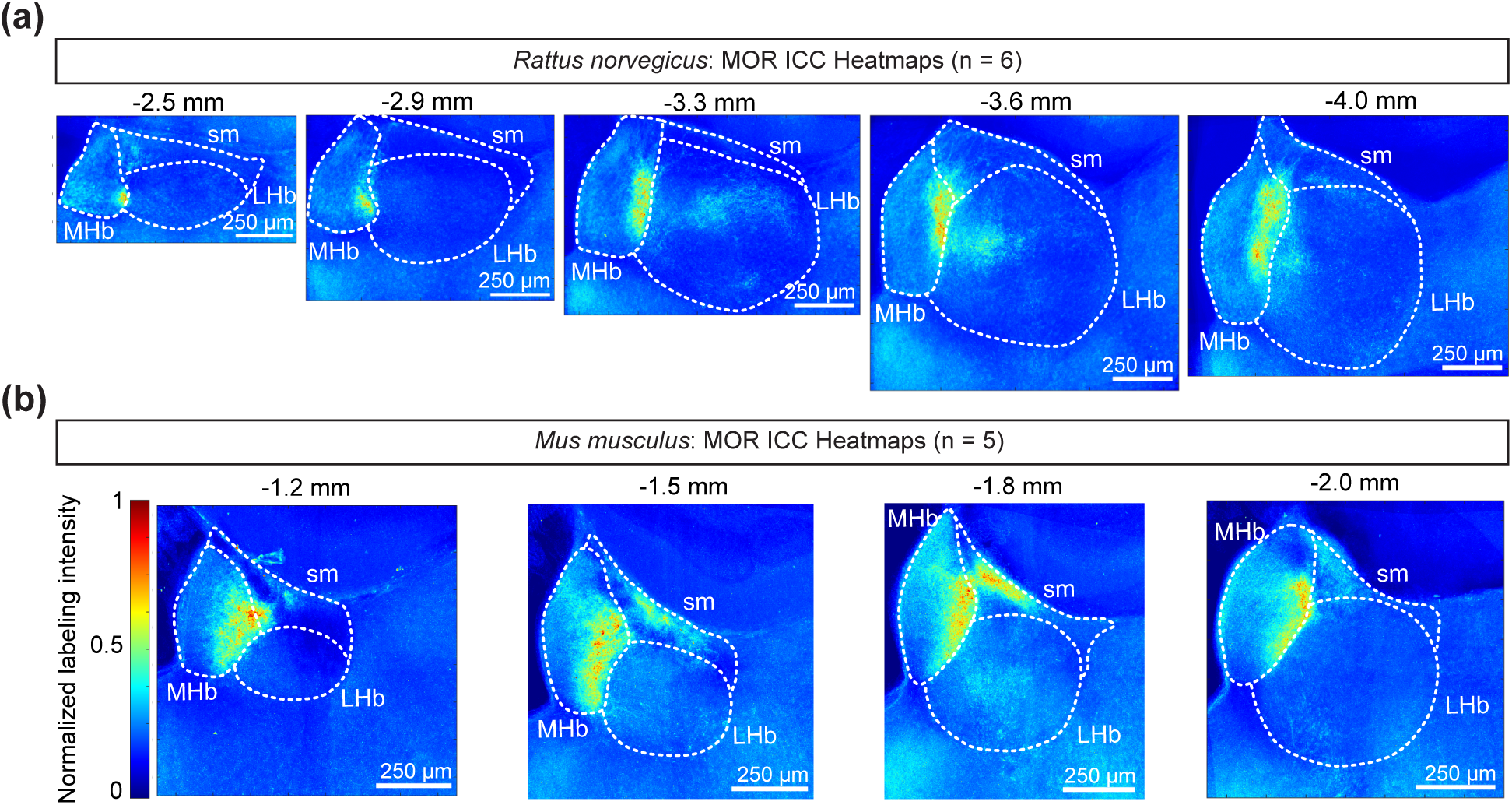
Summary heatmaps for MOR protein expression across rodents. **(a)** Heatmaps of MOR expression across AP levels of the habenula in *Rattus norvegicus*. Heatmaps generated from normalized MOR ICC intensity aligned and averaged across Sprague-Dawley rats (n = 6. **(b)** Heatmaps of MOR expression across AP levels of the habenula in *Mus musculus*. Sections were aligned and MOR ICC averaged across C57Bl6/J mice (n = 5). Scale bars = 250 μm. LHb, lateral habenula; MHb, medial habenula; sm, stria medullaris; fr, fasciculus retroflexus.

### 3.5 MOR-Venus distribution in the *Oprm1-^Venus/Venus^*Hb

Finally, we mapped the signal of fluorescently tagged MORs in *Oprm1-^Venus/Venus^*mice (N = 3 female, 2 male). In these mice fluorescence should indicate the precise location of MOR protein. To amplify the endogenous signal, sections were immunolabeled with an anti-GFP antibody (Figure 6). We used DAPI as a counterstain to delineate the subregions of the Hb complex.

**Figure 6.**
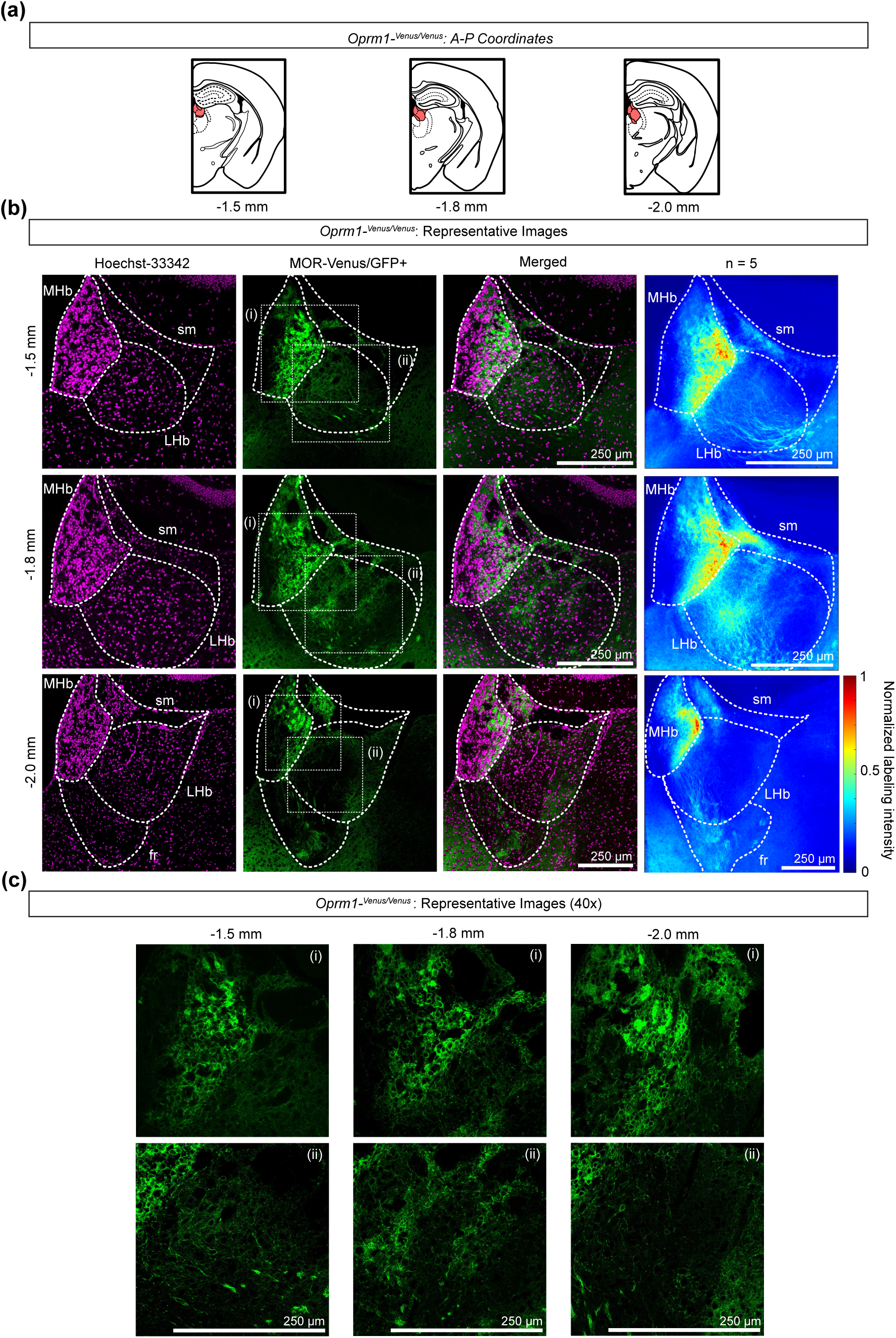
MOR-Venus signal from *Oprm1-^Venus/Venus^* amplified with GFP ICC. **(a)** Depictions of coronal sections at the three AP levels chosen for immunocytochemical processing (∼ -1.5 mm, -1.8 mm, -2.0 mm from bregma). The habenular complex is highlighted in red. Coordinates and drawings are based on the mouse brain atlas (Franklin and Paxinos, 2007). **(b)** Representative images of Hoechst-33342 (magenta), MOR-Venus/GFP ICC (green), and merged signal were chosen from one mouse. (right) Heatmaps of amplified MOR-Venus/GFP signal aligned and summarized across the habenula of *Oprm1-^Venus/Venus^* mice. Heatmaps generated from normalized labeling intensity of MOR-Venus/GFP signal averaged across Hb sections (n = 5). **(c)** Images of MHb (i) and LHb (ii) of the boxed areas in **(b)**. Images in **(b)** and **(c)** depict one focal plane of their respective z-stacks. Scale bars = 250 μm. LHb, lateral habenula; MHb, medial habenula; sm, stria medullaris; fr, fasciculus retroflexus.

We observed extensive MOR-Venus signal in the MHb (Figure 6b). This signal was brightest in the dorsal and lateral portions of the MHb, with little-to-no signal in the medial third (Figure 6b,c). This labeling predominantly outlined cell bodies (Figure 6c). At the most posterior level, this bright signal occupied a smaller proportion of the lateral MHb (Figure 6b).

We also detected MOR-Venus labeling in the LHb at all AP coordinates (Figure 6b,c). At -1.5 mm posterior to bregma there was extensive signal across the LHb that was less intense than in the neighboring MHb (Figure 6b). This included dispersed labeling in fibers medially, and bright labeling of fibers in the ventrolateral LHb (Figure 6b). In intermediate AP sections (-1.8 mm posterior to bregma) we observed higher intensity signal in cell bodies and fibers, including a dense area centered in the medial LHb (Figure 6b). At -2.0 mm posterior to bregma MOR-Venus signal was present at much lower levels although outlined cell bodies and fibers were still clear throughout the LHb (Figure 6b,c). Higher intensity labeling here was localized to medial LHb, along the border with MHb and entering the fr (Figure 6b,c).

Averaged across *Oprm1-^Venus/Venus^* animals we observed bright MOR-Venus signal in the lateral portion of the MHb (Figure 6b). In the LHb, MOR-Venus signal was localized to the medial edge of the LHb at all AP coordinates across animals (Figure 6b). There was also MOR-Venus expression on fibers in the fr, as well as in the sm (Figure 6b). In the LHb, mean MOR-Venus signal was brightest and most extensive in intermediate AP sections (-1.8 mm posterior to bregma) across animals (Figure 6b). There was detectable signal here throughout the majority of the LHb except in the most lateral area. There was a distinct density of MOR-Venus signal at this mid-AP level in the medioventral LHb across animals (Figure 6b).

## 4 Discussion

Here we found *Oprm1* mRNA and MOR protein in both the MHb and LHb with varying distributions across the AP axis in rat and mouse. To accurately delineate MHb from LHb in an unbiased manner, we used cell density, in some cases in combination with *Tac2* or *Tac3* in mouse and rat, respectively. We used methods to detect both mRNA and protein of MOR because the two approaches provide distinct information; mRNA abundance does not necessarily correlate with amount or distribution of translated protein. On average, there was a higher percentage of cells expressing *Oprm1* in the MHb compared to LHb. *Oprm1+* cells in the MHb also typically possessed more puncta per cell than in the LHb. This pattern was consistent across species. As in previous studies of MOR or *Oprm1* distributions, we found high levels of protein and mRNA labeling along the lateral edge of the MHb in both species (Moriwaki et al., 1996; Aizawa et al., 2012; Gardon et al., 2014; Mengaziol et al., 2022). Importantly, we also found MOR and *Oprm1* expression in the mouse and rat LHb at all AP positions. In rat, we found higher-intensity signal for both *Oprm1* and MOR at intermediate AP positions of MHb and LHb. In these sections, cells were labeled for *Oprm1* across the LHb with a greater number of puncta per cell and higher MOR protein expression in the dorsolateral LHb. Additionally, *Oprm1* and MOR were found in the medial LHb in rat, particularly in posterior areas. In the mouse Hb, high-intensity MOR and *Oprm1* signal was detected in MHb throughout the AP axis, while in the LHb intermediate sections possessed a higher percentage of *Oprm1+* cells compared to posterior sections. Averaged MOR signal across animals indicates that there was a higher density of protein localized to the medial LHb. Whether these opioid receptor distribution patterns across species have parallel differences in circuit connectivity and modulation remains to be determined. We also observed some MOR protein distribution divergence between C57Bl6/J mice and a knock-in *Oprm1-^Venus/Venus^*mouse model (see below). That said, overall, *Oprm1* and MOR were clearly detected in both MHb and LHb in both species.

### 4.1 Technical Considerations

In some studies the sm is considered part of the LHb. We observed some *Oprm1* expression in the sm, particularly in mouse, at most levels of the Hb. We decided to exclude this fiber tract from quantification of *Oprm1* because the cells embedded within may be functionally more related to the fibers of passage than the Hb proper.

In the RNAscope analysis we inferred that each puncta represents one probe-bound mRNA molecule within a specific soma, enabling the quantitative interpretation of the number of puncta detected per cell. We counted as “labeled” any cell with two or more puncta because of our very low background noise in this preparation. That said, it is possible that this approach resulted in some type I (false positive) errors. We identified cells with DAPI as it is easily compatible with RNAscope processing, although DAPI labeling does not delineate neurons from non-neuronal cell populations. This technique also does not allow for quantification of mRNA localized to distal dendrites or axons (Dalla Costa et al, 2021).

For immunocytochemistry, the anti-MOR antibody used here targets the C-terminus, intracellular tail of the receptor (AA 384-398). Therefore, the antibody must enter a cell in order to access its binding site on the receptor. It is possible that this led to an underestimation of protein. To be conservative, we do not interpret whether localization is to somatodendritic regions of Hb neurons or to innervating axons, although we did observe labeling that surrounded somata in both the MHb and LHb (Figure 3c, Figure 4c). Overall, these topographical locations were similar to where mRNA was detected.

### 4.2 Topography of mRNA vs protein across species

While the Hb has been shown to have high levels of MOR binding since early autoradiographic studies, the resolution of these methods was not high enough to localize signal to MHb or LHb (Mansour et al., 1987; Quirion et al., 1983; Recht et al., 1985; Herkenham and Pert, 1980).

Previous work using an antiserum against the C-terminus of MOR in rat reported immunoreactivity generally in perikarya in Hb nuclei and in processes in the fr (Moriwaki et al., 1996). They also reported labeling of fibers in the medial portion of the LHb, similar to our MOR expression in the posterior rat Hb, although without an independent counterstain interpretation is difficult (Moriwaki et al., 1996). More recent studies utilizing higher resolution approaches for labeling and detection have proposed that within the Hb most, if not all, MOR expression is found in the MHb (Gardon et al., 2014; Bailly et al., 2020; Wallace et al., 2020; Hashikawa et al., 2020). Our results unequivocally show *Oprm1*+ cells and MOR expression in both the MHb and LHb in mouse and rat. We also mapped the signal in the Hb in a knock-in *Oprm1-^Venus/Venus^*mouse line, an animal designed with the same approach as the MOR-mCherry mouse line (Ehrlich et al., 2019; Erbs et al., 2015; Boulos et al, 2020; Fagan et al., 2024). While overall we found many overlapping patterns of immunocytochemical distributions of MOR in wildtype mice and Venus in *Oprm1-^Venus/Venus^*mice, there were some places of divergence. Most notable are the broader Venus labeling across the MHb and distinct tufts of fibers in the LHb that appear to run into the fr that were not evident in tissue from wildtype animals.

*Oprm1* mRNA mapping reveals the topography of cells that have the ability to translate MOR protein, but does not indicate where the protein is localized and available to respond to ligand binding. *Oprm1* signal was more robust in the mouse Hb compared to rat, both in percentage of *Oprm1* cells and number of puncta per cell, especially in the MHb (Figure 1, Figure 2). *Oprm1* expression in the LHb was quantitatively more similar across species than in the MHb, with high intensity signal in the medial and dorsal LHb of both rat and mouse at intermediate to posterior levels.

MOR protein may be present on or within somata, dendrites, or afferent innervation. Areas with clear immunocytochemical but low *Oprm1* signal could indicate MOR protein localized to afferent inputs. There were several features of overlap between *Oprm1* and MOR protein distributions including in the lateral region of the MHb across the AP axis in rat (Figure 1b, Figure 3b). This pattern was similar but not identical in mouse, where *Oprm1* and MOR were expressed abundantly in a wider area of the lateral MHb (Figure 2b, Figure 4b). This was detected throughout the AP extent of mouse Hb, but varied in detail. In these areas, MORs seemed to localize around somata, but there were many labeled fibers nearby, particularly in the LHb. The physiological responses to MOR activation in the Hb that have been reported include decreasing firing rate of a subset of MHb neurons in mice (Singhal et al., 2025), hyperpolarizing a subset of LHb neurons in rat, and presynaptically inhibiting both glutamate and GABA release onto LHb neurons in rat (Margolis and Fields, 2016). Thus, it is not surprising that we detected MOR protein in these different cellular compartments.

### 4.3 Considerations from gene to circuits

*Oprm1* is a complex gene with 18 exons. Rodents and humans each possess at least 12 splice variants (Shabalina et al., 2009; Liu et al., 2021, Kang et al., 2022), and the RNAscope probe we selected here should detect all *Oprm1* splice variants. These splice variants are differentially distributed across brain regions (Xu et al., 2014), and there is evidence that ligand responses vary across splice variants (Kang et al., 2022). If different splice variants are located in the MHb and LHb, this could enable selective modulation of neural circuits through one or the other Hb region by either endogenous opioid peptides or exogenous drugs.

Recent single cell transcriptomics studies have clustered populations of habenular cells based on genetic profiles. In these studies, the cell cluster with distinct *Oprm1* expression is associated with MHb neurons, although there are groups of LHb neurons with detectable *Oprm1* expression (Wallace et al., 2020; Hashikawa et al., 2020). These studies have also revealed that many *Oprm1* expressing Hb neurons also express the orphan receptor *GPR151*, a provocative pharmacological target since it might function as an “anti-opioid” and its expression is limited to just a few brain regions (Hashikawa et al., 2020; DePasquale et al., 2025). Clusters of MHb neurons with elevated *Oprm1* expression also tend to be distinct from other clusters by also having high expression of *Calb2* and *Cck*. The group of LHb neurons with *Oprm1* expression also express *Lapr1* and *Htr2c* (Hashikawa et al., 2020). Genetic profiling provides unique information about the habenular cells that are capable of producing MORs and may correlate to specific Hb neural connectivity.

The MHb is densely packed with neurons, and most, if not all, of these neurons project to the IPN. Here we found that only the neurons along the lateral border of the MHb showed clear *Oprm1* and MOR expression, and this is consistent with some earlier reports (Aizawa et al., 2012; Gardon et al., 2014; Bailly et al., 2020; Singhal et al., 2025). The IPN has extremely high density of binding and immunocytochemical detection of MOR, at least some of which is localized to MHb axon terminals (Mansour et al., 1987; Kitchen et al., 1997; Bailly et al., 2020). A knock-in mouse line designed to express MOR fused with a red fluorescent protein exhibits strong fluorescent signal in the Hb, all along the fr, and in the IPN (Gardon et al., 2014; Erbs et al., 2015; Bailly et al; 2020). Functional MOR has been demonstrated on the glutamate and glutamate-acetylcholine axon terminals in the IPN arising from the MHb, increasing glutamate release (Singhal et al., 2025), consistent with immunocytochemical localization.

Several studies have used morphology, immunoreactivity, and transcriptomics to divide the LHb into several subregions (Andres et al., 1999; Sylwestrak et al., 2022; Aizawa et al., 2012; Hashiwaka et al, 2020; Iwahori 1977; Behzadi et al., 1990; Root et al., 2015; Skagerberg, 1984; Geisler et al., 2003). Tracing studies often broadly categorize the LHb into medial and lateral subregions (LHbM and LHbL, respectively) as this most closely relates to observed input and output patterns, although even at this level strict compartmentalization has not been reported (Herkenham et al., 1977; Zahm and Root, 2017). We previously reported that glutamatergic innervation of the LHb from the LPO is more strongly inhibited by MOR activation than other glutamatergic inputs (Waung et al., 2022). LPO glutamatergic axons are largely concentrated in the medial aspect of the LHb (Barker et al., 2017), where we also detected higher MOR expression in mid-posterior mouse and rat LHb. Input from the EPN is localized to the lateral portion of the LHb (Wallace et al., 2017; Geisler et al., 2003) where we detected less MOR expression, consistent with our previous functional finding that these inputs are minimally sensitive to MOR activation (Waung et al., 2022).

In greater anatomical detail, Andres et al. (1999) proposed 5 subnuclei within the MHb and 10 subnuclei within the LhbM and LHbL in rat, differentiated by analysis of neuropil at the semithin section level (Figure 7a). Some of these overlap with the sm. As described by Aizawa et al., the superior subnucleus of the MHb (MHbS) contains exclusively glutamatergic neurons, the dorsal region of the central subnucleus of the MHb (MHbC) contains substance P-ergic and glutamatergic neurons, and both cholinergic and glutamatergic neurons are found in the ventral subnuclei of the MHb, including the lateral subnucleus of the MHb (MHbL) where we saw high expression of *Oprm1* and MOR (Aizawa et al., 2012). In the LHb, Zahm and Root (2017) noted that most function and connectivity is broadly distributed rather than confined to any subnucleus, with the exceptions of the central subnucleus of the LHbM (LHbMC) and the parvocellular subnucleus of the LHbM (LHbMPc); there is evidence that neurons in these subnuclei project preferentially to the raphe nuclei and these regions exhibit high immunoreactivity to tyrosine hydroxylase in afferents originating from the VTA (Skagerberg et al., 1984; Zahm and Root, 2017). Using this framework, we detected higher intensity *Oprm1* and MOR labeling in LHbMC and LHbMPc in the intermediate to posterior rat Hb (Figure 7a). We also found high MOR expression in the dorsolateral LHb, overlapping with the LHbLPc. Subnuclei have more recently been defined in the mouse Hb (Figure 7b), and similar to the rat MHb, we saw high intensity labeling of *Oprm1* and MOR in the lateral subnucleus of the ventral division of the mouse MHb (MHbVl) (Wagner et al., 2014). While the subnuclei defined in the mouse Hb largely correspond with rat anatomy, tyrosine hydroxylase immunoreactivity in fibers is notably absent in the mouse LHbMPc (Wagner et al., 2014). Although the pattern of MOR distribution in C57Bl6/J mouse LHb did not strongly correspond to these subnuclei, there was more intense labeling in the ventral part of the LHbLMC in *Oprm1-^Venus/Venus^* mice that did not appear to be fibers of passage.

**Figure 7.**
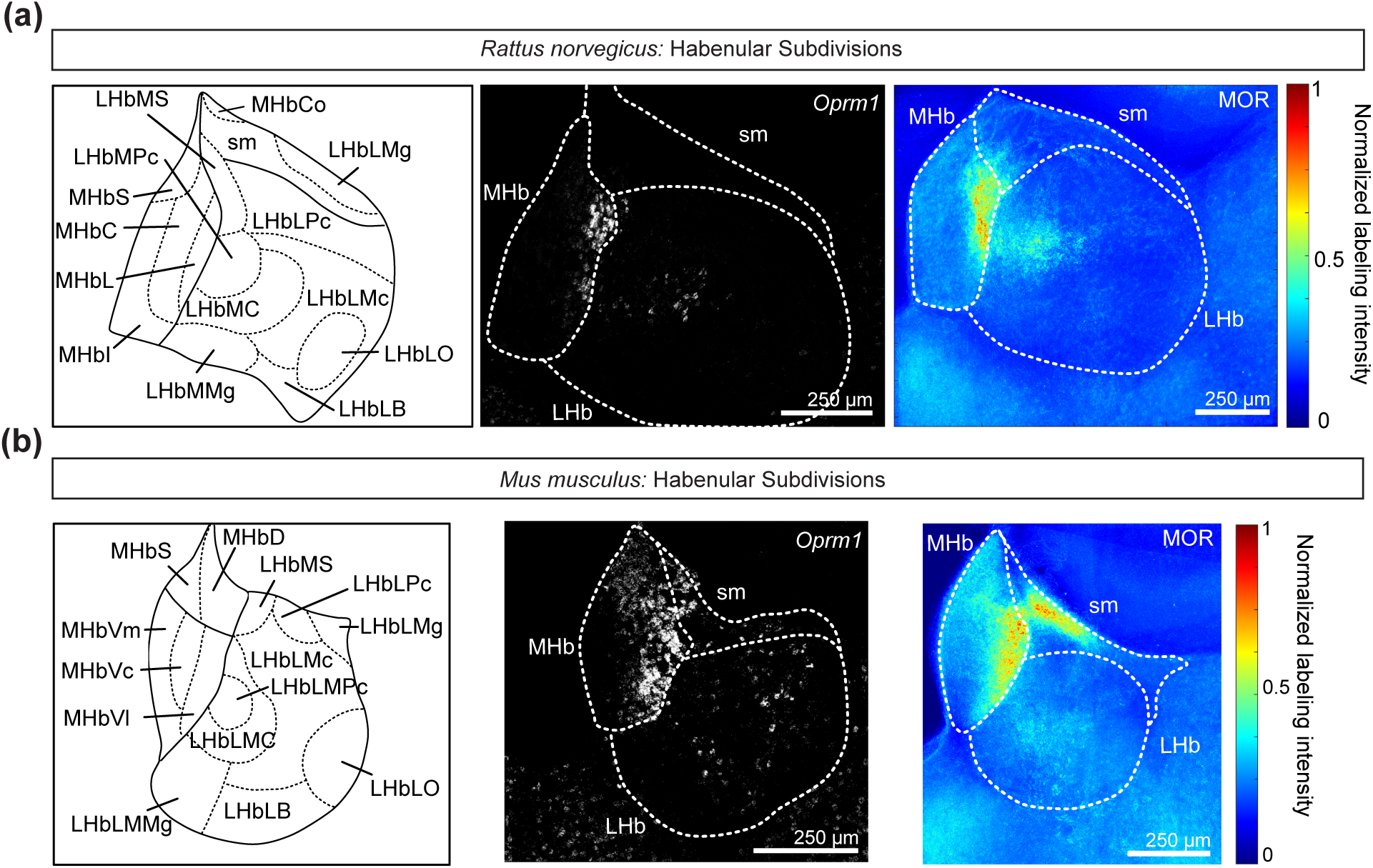
Subdivisions of the MHb and LHb for *Rattus norvegicus* and *Mus musculus*. **(a)** Left, schematic of the posterior rat habenula, with the LHb further divided into eight subsections based on Andres et al. and reviewed by Zahm and Root (Andres et al., 1999; Zahm and Root, 2017). Middle, representative image of *Oprm1* RNAscope in the rat Hb at a similar AP level (∼-4.0 from bregma). Right, summary heatmap of MOR ICC averaged across rats in the posterior habenula (∼-3.6 mm from bregma); panel reproduced from Figure 5a. **(b)** Left, schematic of the posterior mouse habenula, with the LHb further divided into eight subsections based on Wagner et al. (Wagner et al., 2014). Middle, representative image of *Oprm1* RNAscope in the mouse Hb at a similar AP level (∼-1.8 from bregma). Right, heatmap of MOR ICC averaged across mice in the posterior habenula (∼-1.8 mm from bregma); panel reproduced from Figure 5b. Scale bars = 250 μm. LHb, lateral habenula; LHbLB, basal subnucleus of the lateral division of the LHb; LHbLMc, magnocellular subnucleus of the lateral division of the LHb; LHbLMg, marginal subnucleus of the lateral division of the LHb; LHbLO, oval subnucleus of the lateral division of the LHb; LHbLPc, parvocellular subnucleus of the lateral division of the LHb; LHbMC, central subnucleus of the medial division of the LHb; LHbMMg, marginal subnucleus of the medial division of the LHb; LHbMPc, parvocellular subnucleus of the medial division of the LHb; LHbMS, superior subnucleus of the medial division of the LHb; MHb, medial habenula; MHbC, central part of the MHb; MHbCo, commissural part of the MHb; MHbD, dorsal subnucleus of the MHb; MHbI, inferior part of the MHb; MHbL, lateral part of the MHb; MHbS, superior part of the MHb; MHbVc, central subnucleus of the ventral division of the MHb; MHbVl, lateral subnucleus of the ventral division of the MHb; MHbVm, medial subnucleus of the ventral division of the MHb; sm, stria medullaris.

We did not observe clear sex differences in anatomical MOR distributions in the Hb, thus all analyses are presented as averages across sexes. However, some sex differences in MOR function have been reported for the Hb at the level of behavior. We previously showed that MOR activation inhibits glutamate release from LPO axon terminals onto LHb neurons similarly in male and female rats (Waung et. al., 2022). Yet behaviorally we found that microinjection of a MOR agonist into the LHb produced both sensory and affective relief of ongoing pain in male rats, but only affective pain relief in females (Waung et. al., 2022). Females also required a higher dose of MOR agonist for this effect (Waung et al., 2022). Since we did not detect clearsex differences in anatomical labeling of MORs in this study or in MOR modulation of synaptic responses (Waung et al., 2022), it is possible that the behavioral sex differences we observed were due to circuit level neural connectivity differences between males and females.

## Conclusion

Together these observations locate *Oprm1* mRNA and MOR protein in both MHb and LHb in rat and mouse. This work complements *ex vivo* physiological evidence for functional MOR expression in both regions (Singhal et al, 2025; Margolis and Fields, 2016; Waung et al., 2022). The fact that the input and output connections of the MHb and LHb are essentially non-overlapping means that the functions of these two MOR controlled brain regions should be considered to be independent. Understanding the anatomical distributions and physiological impact(s) of MOR activation combined with circuit-based behavioral studies of these two neighboring brain regions will enable improved understanding of how MOR activation contributes to mood and behavior.

## Acknowledgments

We thank Ben Ngu and Alex J. Keip for their technical assistance. This work was supported by National Institutes of Health grants R01NS134981 and R01DA042025 to EBM; National Institutes of Health grants 5R01DA059388-03 and 5R37MH119156-06 to CCH and RCS; National Institute of Mental Health grant K01MH123757 to ATE; A.P. Giannini Foundation support to RCS; Coordenação de Aperfeiçoamento de Pessoal de Nível Superior grant 88887.878571/2023-00 to BBA; funds from the Painless Research Foundation; and funds from the State of California for medical research on alcohol and substance use.

## Data Availability

All raw data are available on the Brain Image Library at DOI: https://doi.org/10.35077/g.1205

