## Supplemental Table 1 for "Mu opioid receptor mRNA and protein localization across the rat and mouse habenula"

**Supplementary Table 1:** Statistical testing for summary data

| Experiment | Figure | Test Used | Statistics (test statistic; p-value) | Multiple Comparison Correction (Test Used): Mean, C.I., and p-value |
|---|---|---|---|---|
| Percentage of *Oprm1+* cells, MHb vs LHb (rat) | 1c | Unpaired t-test, two-tailed | t = 8.748, df = 18; p < 0.0001 | n/a |
| Percentage of *Oprm1+* cells across AP positions and Hb subregion (rat) | 1c | Mixed effects modeling (Type III ANOVA) | AP position: F(1.779, 16.01) = 9.533; p = 0.0024<br><br>Hb subregion: F(1.000, 9.000) = 147.7; p < 0.0001<br><br>AP position x Hb subregion: F(1.822, 7.895) = 9.136; p = 0.0097 | Pairwise comparisons (Tukey's HSD):<br><br>MHb:<br><br>-2.5 mm vs -3.0 mm: -12.06, [-22.78, -1.335]; p = 0.0287<br><br>-2.5 mm vs -3.5 mm: -4.11, [-13.10, 4.876]; p = 0.4984<br><br>-2.5 mm vs -4.0 mm: -11.11, [-47.57, 25.35]; p = 0.3784<br><br>-3.0 mm vs -3.5 mm: 7.356, [0.6150, 14.10]; p = 0.0326<br><br>-3.0 mm vs -4.0 mm: 1.091, [-34.46, 36.64]; p = 0.9986<br><br>-3.5 mm vs -4.0 mm: -4.947, [-33.78, 23.88]; p = 0.8397<br><br>LHb:<br><br>-2.5 mm vs -3.0 mm: -16.39, [-28.12, -4.656]; p = 0.0089<br><br>-2.5 mm vs -3.5 mm: 3.47, [-7.015, 13.95]; p = 0.7215<br><br>-2.5 mm vs -4.0 mm: 1.678, [-7.755, 11.11]; p = 0.6687<br><br>-3.0 mm vs -3.5 mm: 18.4, [10.02, 26.79]; p = 0.0004 |

| | | | | |
|---|---|---|---|---|
| | | | | -3.0 mm vs -4.0 mm: 12.63, [-30.29, 55.55]; p = 0.5661<br><br>-3.5 mm vs -4.0 mm: -3.008, [-30.51, 24.49]; p = 0.9467 |
| Number of puncta per *Oprm1*+ cell, MHb vs LHb (rat) | 1d | Welch's t-test, two-tailed | t = 8.748, df = 18; p < 0.0001 | n/a |
| Number of puncta per *Oprm1*+ cell across AP positions and Hb subregion (rat) | 1d | Mixed effects modeling (Type III ANOVA) | AP position: F(1.715, 15.43) = 11.76; p = 0.0011<br><br>Hb subregion: F(1.000, 9.000) = 70.49; p < 0.0001<br><br>AP position x Hb subregion: F(1.846, 7.999) = 4.384; p = 0.0539 | Pairwise comparisons (Tukey's HSD):<br><br>MHb:<br><br>-2.5 mm vs -3.0 mm: -7.427, [-13.30, -1.554]; p = 0.0156<br><br>-2.5 mm vs -3.5 mm: -7.238, [-14.29, -0.1827]; p = 0.0445<br><br>-2.5 mm vs -4.0 mm: -10.93, [-25.72, 3.862]; p = 0.1077<br><br>-3.0 mm vs -3.5 mm: 0.2548, [-3.800, 4.310]; p = 0.9971<br><br>-3.0 mm vs -4.0 mm: -3.997, [-17.08, 9.085]; p = 0.5425<br><br>-3.5 mm vs -4.0 mm: -5.409, [-24.60, 13.78]; p = 0.5927<br><br>LHb:<br><br>-2.5 mm vs -3.0 mm: -2.161, [-4.154, -0.1675]; p = 0.0343 |

| | | | | |
|---|---|---|---|---|
| | | | | -2.5 mm vs -3.5 mm: -2.81, [-4.483, -1.136]; p = 0.003<br><br>-2.5 mm vs -4.0 mm: -3.276, [-7.025, 0.4734]; p = 0.071<br><br>-3.0 mm vs -3.5 mm: -0.6609, [-2.681, 1.359]; p = 0.7418<br><br>-3.0 mm vs -4.0 mm: -1.044, [-6.289, 4.200]; p = 0.7801<br><br>-3.5 mm vs -4.0 mm: -0.2782, [-2.986, 2.430]; p = 0.9549 |
| Percentage of *Oprm1*+ cells, MHb vs LHb (mouse) | 2c | Unpaired t-test, two-tailed | t = 7.101, df = 22; p < 0.0001 | n/a |
| Percentage of *Oprm1*+ cells across AP positions and Hb subregion (mouse) | 2c | Mixed effects modeling (Type III ANOVA) | AP position: F(1.395, 15.35) = 3.294; p = 0.0783<br><br>Hb subregion: F(1.000, 11.000) = 178.1; p < 0.0001<br><br>AP position x Hb subregion: F(1.554, 8.807) = 6.757; p = 0.0207 | Pairwise comparisons (Tukey's HSD):<br><br>MHb:<br><br>-1.2 mm vs -1.5 mm: -6.089, [-19.77, 7.595]; p = 0.5356<br><br>-1.2 mm vs -1.8 mm: -2.202, [-19.04, 14.64]; p = 0.9757<br><br>-1.2 mm vs -2.0 mm: -1.134, [-16.65, 14.38]; p = 0.9937<br><br>-1.5 mm vs -1.8 mm: 3.055, [-5.606, 11.72]; p = 0.6978<br><br>-1.5 mm vs -2.0 mm: 7.241, [-3.727, 18.21]; p = 0.2034<br><br>-1.8 mm vs -2 mm: 5.395, [-12.92, 23.71]; p = 0.7119 |

| | | | | |
|---|---|---|---|---|
| | | | | LHb:<br><br>-1.2 mm vs -1.5 mm: -11.39, [-39.22, 16.44]; p = 0.5979<br><br>-1.2 mm vs -1.8 mm: 6.649, [-19.91, 33.21]; p = 0.861<br><br>-1.2 mm vs -2.0 mm: 13.15, [-7.535, 33.83]; p = 0.2251<br><br>-1.5 mm vs -1.8 mm: 15.4, [5.159, 25.63]; p = 0.0051<br><br>-1.5 mm vs -2.0 mm: 24.99, [6.189, 43.78]; p = 0.0145<br><br>-1.8 mm vs -2 mm: 12.38, [-12.43, 37.18]; p = 0.3556 |
| Number of puncta per *Oprm1*+ cell, MHb vs LHb (mouse) | 2d | Welch's t-test, two-tailed | t = 7.101, df = 22; p < 0.0001 | n/a |
| Number of puncta per *Oprm1*+ cell across AP positions and Hb subregion (mouse) | 2d | Mixed effects modeling (Type III ANOVA) | AP position: F(1.148, 12.63) = 1.957<br><br>Hb subregion: F(1.000, 11.000) = 64.45; p < 0.0001<br><br>AP position x Hb subregion: F(1.925, 10.91) = 2.173; p = 0.1615 | n/a |
