## Supplemental Figure 1 for "Mu opioid receptor mRNA and protein localization across the rat and mouse habenula"

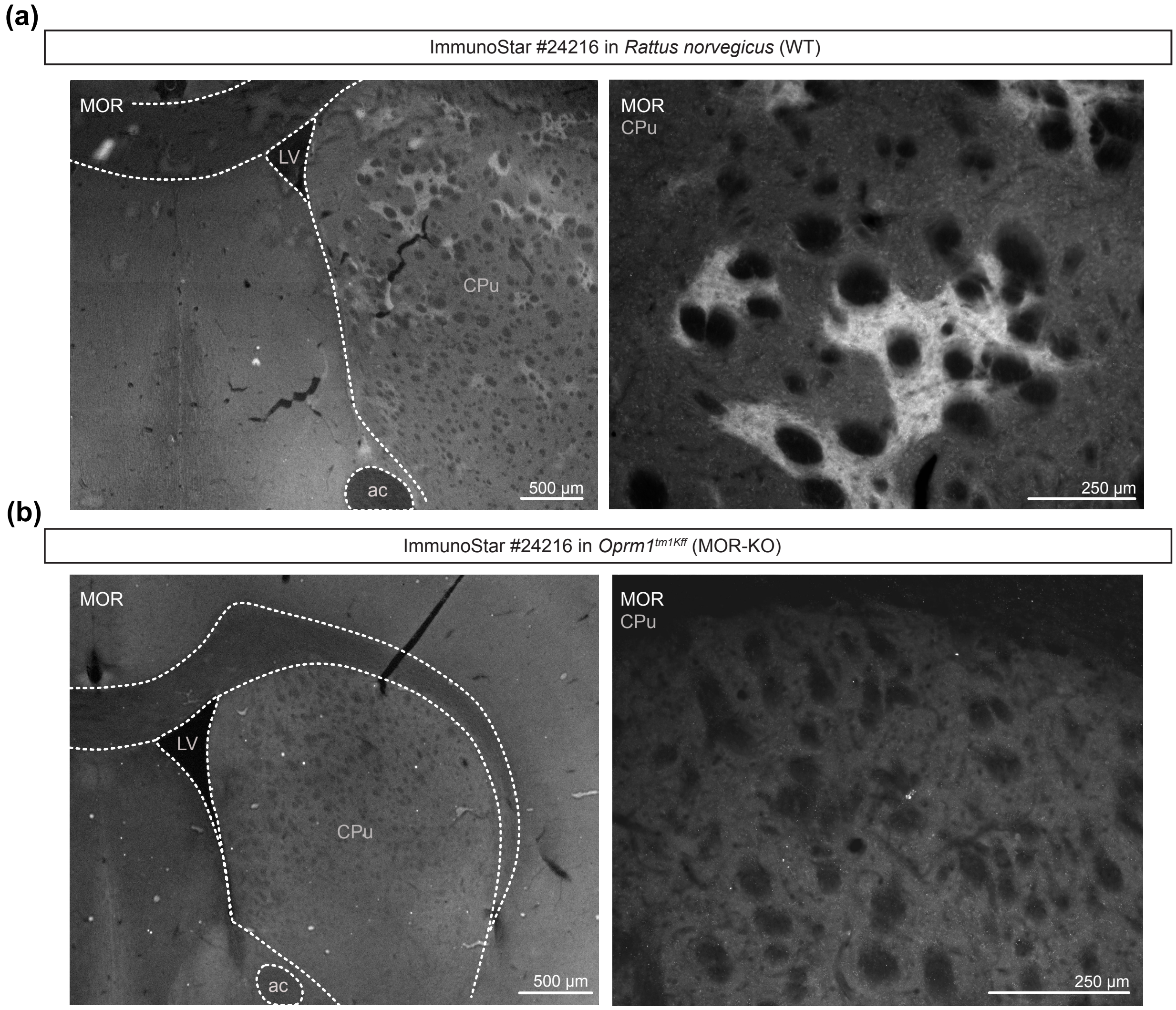


**Supporting Information Figure 1.** Antibody specificity of ImmunoStar #24216 in tissue from MOR-KO (*Oprm1*$^{tm1Kff}$) mouse and WT rat. **(a)** Immunocytochemistry for MOR protein in the dorsal striatum of a WT rat. MOR signal in the CPu captured at low magnification (left, scale bar = 500 µm) and higher magnification (right, scale bar = 250 µm). **(b)** Immunocytochemistry for MOR protein in the dorsal striatum of a MOR-KO mouse (*Oprm1*$^{tm1Kff}$*,* RRID:IMSR_JAX:007559). MOR signal in the CPu captured at low magnification (left, scale bar = 500 µm) and higher magnification (right, scale bar = 250 µm). ac, anterior commissure; CPu, caudate-putamen; LV, lateral ventricle.
