## Supplemental Figure 2 for "Mu opioid receptor mRNA and protein localization across the rat and mouse habenula"

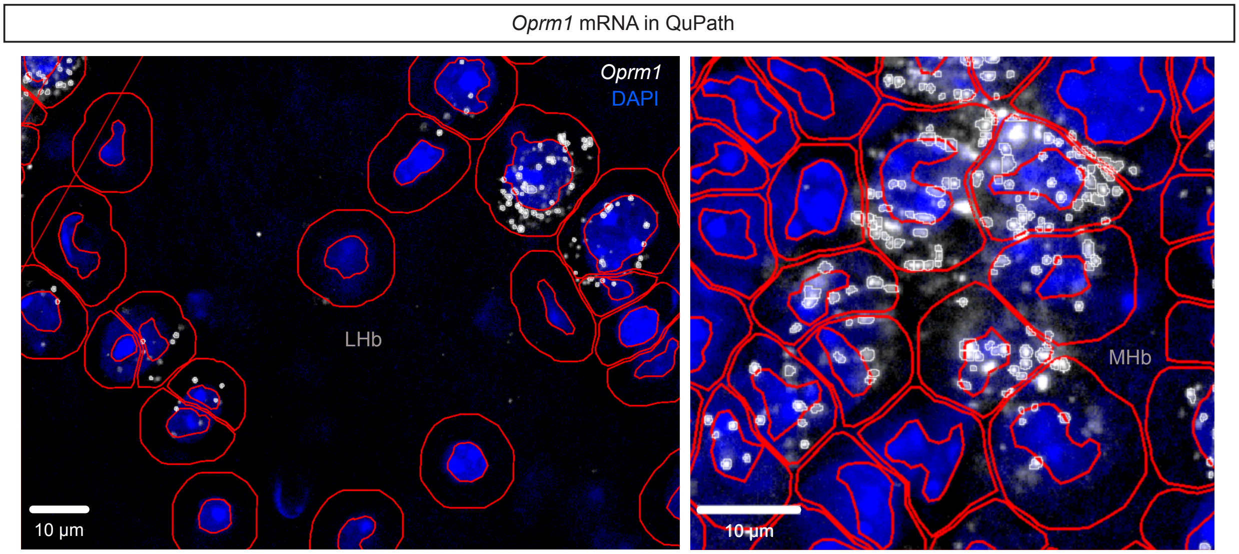


**Supporting Information Figure 2.** Example images of low density (LHb, left) and high density (MHb, right) *Oprm1* mRNA puncta in QuPath. DAPI (blue) was used for nucleus detection and cell body estimation (red outlines), and *Oprm1* mRNA (white) was detected using subcellular detection (white outlines). Scale bars = 10 µm. LHb, lateral habenula; MHb, medial habenula.
